# On testing structural identifiability by a simple scaling method: relying on scaling symmetries can be misleading

**DOI:** 10.1101/2020.11.30.403956

**Authors:** Alejandro F. Villaverde, Gemma Massonis Feixas

## Abstract

A recent paper (Castro M, de Boer RJ, “Testing structural identifiability by a simple scaling method”, PLOS Computational Biology, 2020, 16(11):e1008248) introduces the Scaling Invariance Method (SIM) for analysing structural local identifiability and observability. These two properties define mathematically the possibility of determining the values of the parameters (identifiability) and states (observability) of a dynamic model by observing its output. In this note we warn that SIM considers scaling symmetries as the only possible cause of non-identifiability and non-observability. We show that other types of symmetries can cause the same problems without being detected by SIM, and that in those cases the method may yield a wrong result. Finally, we demonstrate how to analyse structural local identifiability and observability with symbolic computation tools that do not exhibit those issues.

---

The existence of symmetries in the equations of a dynamic model is a source of lack of structural identifiability and observability. Such symmetries [1,2] allow for similarity transformations [3], that is, transformations of parameters and state variables that leave the model output invariant [4]. Since the parameters and states involved in such symmetries cannot be distinguished by observing the output, they are unidentifiable and unobservable. The SIM test proposed in [5] adopts this approach. It starts by decomposing the dynamic equations of a model as a sum of functionally independent functions. Next, unknown parameters and unobserved states are multiplied by unknown scaling factors, and each functionally independent function is equated to its scaled version. Finally, combinations of the scaling factors that leave the equations invariant are sought. The SIM classifies the parameters (respectively, state variables) with a scaling factor equal to one as identifiable (respectively, observable).

Thus, the SIM approach to analysing structural identifiability and observability is to search for a particular type of symmetry, namely scaling symmetries. However, other types of symmetries can also be present in ordinary differential equation (ODE) models. A number of examples from biology have been discussed in the recent literature, see e.g. [6–8]. If the equations of a model only have symmetries that are not of the scaling type, the SIM test does not detect them and wrongly classifies the related parameters as structurally identifiable and the related state variables as observable.

SIM’s limitation to scaling symmetries is mentioned in [5] (“our identifiability test (...) provides a simple way to find a type of symmetry that is related to scale invariance”). However, that paper does not mention that due to this limitation the SIM test can yield wrong results; instead, it claims that “scaling invariance of the model equations can be used to determine whether the parameters are unidentifiable or not”. Indeed, the existence of a scaling invariance indicates that the parameters are unidentifiable. However, the opposite is not true, i.e. its absence does not mean that the parameters are identifiable. We discuss two counter-examples to illustrate this risk.

## Counter-example 1: the FitzHugh-Nagumo model

We first consider the classical FitzHugh-Nagumo model, whose structural identifiability and symmetries were discussed in [6]. It is a nonlinear model that can describe an excitable system such as a neuron, and it can exhibit oscillatory behaviour. Its equations are:

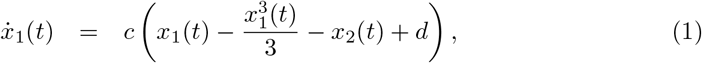

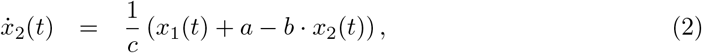

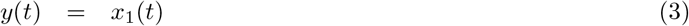

where the *x_i_*(*t*) are the states, *y*(*t*) is the measurable output, and *a, b, c, d* are unknown parameters. In what follows we omit the dependency on *t* to simplify the notation.

We show the calculations of the SIM test below. Briefly, each functionally independent term in (1) and (2) is equated to its scaled counterpart, which introduces scaling factors u_*_ for every state and parameter (except for *x*_1_, which is directly measured). The scaled terms in the ODE of 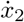 are divided by *u*_*x*_2__ to account for the fact that the derivative of the state is also scaled. More details about the method can be found in [5]. The procedure yields the following equations, where (4)-(6) come from (1) and (7)-(8) come from (2):

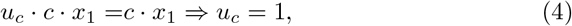

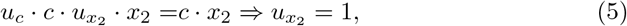

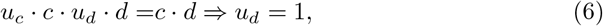

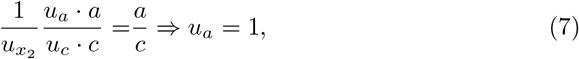

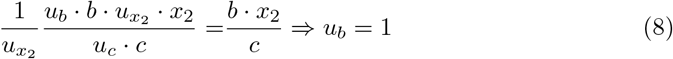

Since the only possible solution is the trivial one (i.e. all the scaling factors are equal to one), the SIM test classifies this model as structurally locally identifiable (s.l.i.) and observable.

However, this result is incorrect: the model is in fact unidentifiable and unobservable, due to the existence of an affine symmetry [6]. This result, which we obtained with the STRIKE-GOLDD toolbox [9], can be easily verified analytically, as we now show. The symmetry analysis, performed using the procedure described in [8], finds the following symmetries, expressed as one-parameter Lie groups of transformations:

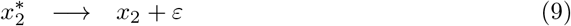

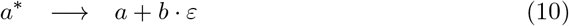

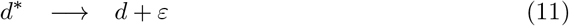

The symmetries are defined as a function of a new parameter *ε*. The above expressions (9)-(11) indicate that replacing the terms to the left of the arrow with their right hand equivalents does not modify the model output. Replacing the right hand expressions of (9)-(11) in the model ODEs (1)-(2) it is immediate to see that the above transformations leave the model equations invariant:

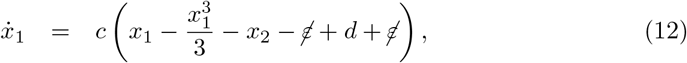

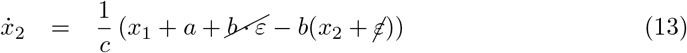

Thus, the state *x*_2_ and the parameters *a, d* cannot be distinguished from their transformed values (9)–(11), and are therefore unobservable and structurally unidentifiable, respectively. However, since the cause of this lack of structural identifiability and observability is not a scaling symmetry but an affine one, SIM wrongly classifies them as observable and structurally identifiable.

## Counter-example 2: a linear compartment model

This second case study shows that not only nonlinear models can have non-scaling symmetries. The following linear compartment model was presented as Example 6.3 in [10], where it was reported that it is unidentifiable and has no identifiable scaling reparameterization. The model consists of four states (one of which is directly measured, *x*_1_), ten parameters, and one known input, *g*:

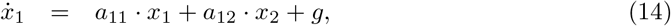

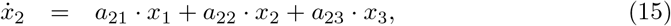

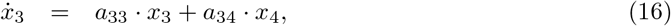

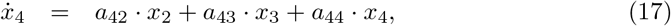

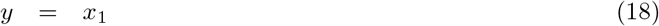

The model above is structurally unidentifiable and non-observable due to the existence of scaling symmetries and one higher order Lie symmetry. Reformulating the model by removing the scaling symmetries, it is possible to obtain an equivalent model with only seven parameters:

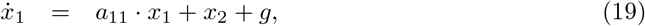

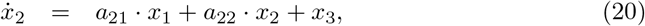

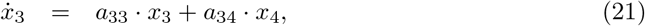

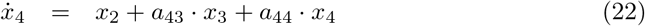

where the measured output is again *y* = *x*_1_. It can be readily seen, following the same procedure as in the previous example, that the SIM test classifies this model as structurally locally identifiable (s.l.i.) and observable. The details of the calculations are not shown here for brevity.

However, this result is incorrect: in fact, only *a*_11_, *a*_21_, *a*_22_ and *a*_34_ are s.l.i.. The remaining parameters (*a*_33_, *a*_43_ and *a*_44_) are structurally unidentifiable, and *x*_4_ is non-observable. As in the previous example, we can verify the correctness of this result (obtained with the STRIKE-GOLDD toolbox) by examining the symmetries of the model, which can be written as follows:

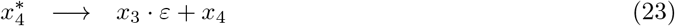

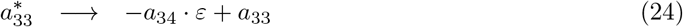

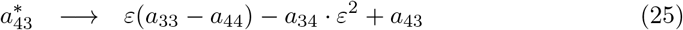

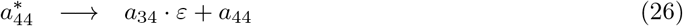

Following the same procedure as in the previous example, it can be checked that, by replacing in the model equations (21) and (22) the left hand side of (23)–(26) with the right hand side, the transformations are cancelled and we obtain the original equations again (calculations not shown). This proves that the model is structurally unidentifiable, and therefore the result of the SIM test is incorrect.

## Computational tools can assess structural identifiability reliably and efficiently

The previous example demonstrates that even relatively simple models can have symmetries that make them unidentifiable and non-observable, and which are not of the scaling type. The counter-examples have also shown that, while the SIM test does not find these symmetries, it is possible to analyse the structural identifiability and observability of these models with symbolic computation methods. We note that the computational cost of these analyses is fairly low: the run-times of the structural identifiability analyses of the first and second counter-examples were 3 and 25 seconds in STRIKE-GOLDD, respectively. When we double-checked the results using the COMBOS web app [11] the run-times were even lower (< 1 and 7 seconds, respectively), although in fairness it should be noted that COMBOS does not analyse observability. The time required for entering the models was, at most, a few minutes in each tool.

These results prompt us to comment on another aspect of [5], namely its assessment of the performance of computational methods. That paper analyses thirteen models with a number of symbolic computation methods, which, when compared to SIM, are portrayed as less applicable, less conclusive, and/or producing incompatible results. However, we have found that the performance of at least some of those computational tools is misrepresented. In particular, we have examined the case of the STRIKE-GOLDD toolbox. In [5] it is claimed that the STRIKE-GOLDD toolbox cannot analyse four of the models (HIV (2), Glycolysis, High dimensional model, and Within-host virus model) and yields wrong results for a fifth one (NF*κ*B (2), which it allegedly classifies as unidentifiable when it is structurally identifiable). In contrast, we analysed these five models with STRIKE-GOLDD and obtained conclusive results in all cases. We summarize the results in Table 1. We speculate that the issues reported in [5] could be due to incorrect specifications of the model definitions (e.g. using reserved variable names) or options (e.g. wrong number of inputs). To clarify this point we provide STRIKE-GOLDD implementations of the five models listed in Table 1, as well as their corresponding options files, as supplementary information.

**Table 1.** Structural identifiability and observability analysis results obtained with STRIKE-GOLDD.

| Model | Result | Run-time (s) | Agrees with SIM |
| --- | --- | --- | --- |
| HIV (2) | s.l.i., observable | 0.6 | yes |
| Glycolysis | s.l.i., observable | 1.3 | yes |
| High dimensional | s.l.i., observable | 1.6 | yes |
| NFκB (2) | s.l.i., unobservable ($x_{15}$) | 2218 | yes* |
| Within-host virus | s.l.i., observable | 8.9 | yes |

Results of the structural identifiability and observability analysis of five models, obtained with STRIKE-GOLDD; s.l.i stands for structurally locally identifiable. In [5] it is claimed that STRIKE-GOLDD yields wrong results about NF*κ*B (2) and cannot analyse the remaining models. Using STRIKE-GOLDD we obtained that NF*κ*B (2) is s.l.i. and one of its states (*x*_15_) is not observable. The latter result is in disagreement with [5]; however, since *x*_15_ does not appear in any equation of the model other than its own, it is not measured, and its initial condition is unknown, it cannot be observable even if all parameters are known [12]. We would also like to note that the run-time for NF*κ*B (2) is much longer than for the other models. This is due to the way in which STRIKE-GOLDD finds the non-observable state, which entails recalculating the observability rank for each model variable. This is a particularly unfavourable case, but the run-time is still in the order of minutes.

These results demonstrate that there is at least one computational method, STRIKE-GOLDD, that provides conclusive and correct results for all the models analysed in [5], as well as for the two case studies used in the present paper. Having said that, we remark that it is not our purpose to advocate the use of STRIKE-GOLDD as the ultimate tool. Rather, our claim is that symbolic computing methods are nowadays a mature solution for performing structural identifiability analyses, although naturally every method has limitations. Besides STRIKE-GOLDD, a number of software tools such as SIAN [13], ObservabilityTest [14], COMBOS [11], EAR [15], GenSSI [16], and DAISY [17], may also be used.

## Simple methods are appealing, but their applicability must be examined with caution

The ability to perform calculations by hand is a desirable feature, not only due to the convenience of not requiring a computing environment, but also because this process can provide unique insights about a problem. In this regard, the SIM test proposed in [5] is appealing and, indeed, it yields correct results in many cases. Unfortunately, it also gives wrong results in other cases, without providing any hint whatsoever. As this note has shown, even apparently simple models can have non-scaling symmetries for which SIM fails. Structural identifiability and observability are properties that often defy intuition, and the search for a simple approach to analyse them has proven elusive for decades. Their analysis usually entails complex symbolic computations that require specialized software. Fortunately, there is a number of well-established tools, available in a variety of computing environments, which can help in this endeavour.

