## Supplementary material for "On testing structural identifiability by a simple scaling method: relying on scaling symmetries can be misleading": options.m

```
%==========================================================================
% THE USER CAN DEFINE THE PROBLEM AND SET OPTIONS IN THE FOLLOWING LINES:                                                       
%==========================================================================

function [modelname,paths,opts,submodels,prev_ident_pars] = options()

%%% (1) MODEL: 
modelname ='FitzHughNagumo'; 

%%% (2) PATHS:
paths.meigo     = '/.../MEIGO';      
paths.models    = strcat(pwd,filesep,'models');
paths.results   = strcat(pwd,filesep,'results');
paths.functions = strcat(pwd,filesep,'functions');
                            
%%% (3) IDENTIFIABILITY OPTIONS:
opts.numeric    = 0;       % calculate rank numerically (= 1) or symbolically (= 0)
opts.replaceICs = 0;       % replace states with known initial conditions (= 1) or use generic values (= 0) when calculating rank
opts.checkObser = 1;       % check state observability, i.e. identifiability of initial conditions (1 = yes; 0 = no).
opts.checkObsIn = 1;       % check input observability (1 = yes; 0 = no).
opts.findcombos = 0;       % try to find identifiable combinations? (1 = yes; 0 = no).
opts.unidentif  = 0;       % use method to try to establish unidentifiability instead of identifiability, when using decomposition. 
opts.forcedecomp= 0;       % always decompose model (1 = yes; 0 = no).
opts.decomp     = 0;       % decompose model if the whole model is too large (1 = yes; 0 = no: instead, calculate rank with few Lie derivatives).
opts.decomp_user= 0;       % when decomposing model, use submodels specified by the user (= 1) or found by optimization (= 0). 
opts.maxLietime = 100;     % max. time allowed for calculating 1 Lie derivative.
opts.maxOpttime = 30;      % max. time allowed for every optimization (if optimization-based decomposition is used).
opts.maxstates  = 6;       % max. number of states in the submodels (if optimization-based decomposition is used).
opts.nnzDerU    = [inf inf inf inf];       % numbers of nonzero derivatives of the measured inputs (u); may be 'inf'
opts.nnzDerW    = 1;       % numbers of nonzero derivatives of the unmeasured inputs (w); may be 'inf'
opts.nnzDerIn   = opts.nnzDerU; % deprecated option

%%% (4) AFFINE OPTIONS:
opts.affine                = 0;     % use algorithm for affine control systems (=1) or not(=0).
opts.affine_tStage         = 1000;  % max. computation time for the last iteration.
opts.affine_kmax           = 4;     % max. number of iterations.
opts.affine_parallel       = 0;     % use parallel toolbox (=1) or not (=0) to calculate partial ranks.
opts.affine_workers        = 4;     % number of workers for parallel pool.
opts.affine_graphics       = 1;     % display graphics (=1) or not (=0)
opts.affine_delete_model   = 1;     % delete affine_model.mat at the end (=1) or not (=0).

%%% (5) User-defined submodels for decomposition (may be left = []): 
submodels = []; 
%- Submodels are specified as a vector of states, as e.g.:
%         submodels{1}  = [1 2];
%         submodels{2}  = [1 3];

%%% (6) MULTI-EXPERIMENT OPTIONS:
opts.multiexp              = 0;    % Execute multi-experiment analysis (=1) or not (=0).
opts.multiexp_numexp       = 3;    % Number of experiments.
opts.multiexp_user_nnzDerU = 0;    % Set manually the number of nonzero derivatives of the measured inputs (u) for each experiment (=1) or not (=0).
opts.multiexp_user_nnzDerW = 0;    % Set manually the number of nonzero derivatives of the unmeasured inputs (w) for each experiment (=1) or not (=0).
opts.multiexp_nnzDerU      = [0 0 0]; % Number of nonzero derivatives of the measured inputs (u) for each experiment; may be 'inf'.
opts.multiexp_nnzDerW      = [0 0 0]; % Number of nonzero derivatives of the unmeasured inputs (w) for each experiment; may be 'inf'.
opts.multiexp_user_ics  = 0;       % Set manually the initial conditions for each experiment (=1) or not (=0).
%- Multi-experiment initial conditions (Rows=variables;Columns:experiments):
opts.multiexp_ics       = [];
% syms x10_1 x10_2 x10_3 x20_1 x20_2 x20_3 x30_1 x30_2 x30_3 
% opts.multiexp_ics     = [x10_1 x10_2 x10_3; x20_1 x20_2 x30_3; x30_1 x30_2 x30_3];
%- Whose initial conditions are known (Rows=variables;Columns:experiments):
opts.multiexp_known_ics = [];
% opts.multiexp_known_ics = [0 1 0;1 0 0;0 1 1];

%%% (7) Parameters already classified as identifiable may be entered below.
prev_ident_pars = [];
%- example:
% syms p2 p5
% prev_ident_pars = [p2 p5];

end
```
