## Supplementary material for "On testing structural identifiability by a simple scaling method: relying on scaling symmetries can be misleading": README.txt

This folder contains Matlab files accompanying the paper "On testing structural identifiability by a simple scaling method: relying on scaling symmetries can be misleading", by Alejandro F. Villaverde and Gemma Massonis Feixas. The files allow to analyse the structural identifiability and observability of the models described in the paper, using the STRIKE-GOLDD toolbox (https://github.com/afvillaverde/strike-goldd).

Instructions for analysing the models:
1. Download the STRIKE-GOLDD toolbox from https://github.com/afvillaverde/strike-goldd
2. Unzip it and copy the files provided here in the 'models' folder, with the exception of 'options.m'.
3. Replace the 'options.m' file included in STRIKE-GOLDD with the one provided here.
4. Select the model to analyse by editing line 8 in the options file.
5. Run 'install.m' from STRIKE-GOLDD's main folder.
6. Run 'STRIKE_GOLDD.m'' from STRIKE-GOLDD's main folder.
 
